# A heterogeneous biomedical knowledge network framework for rare disease drug candidate prioritization: integrating Orphadata and DisGeNET via gene-bridge harmonization

**DOI:** 10.64898/2026.06.16.732556

**Authors:** Diya Ramani

## Abstract

Rare diseases collectively affect over 300 million individuals worldwide, yet the vast majority lack approved pharmacological treatments, leaving patients with few therapeutic options and researchers with limited computational tools for systematic candidate identification. This study presents a network-based computational framework that integrates two complementary public biomedical databases (Orphadata, which catalogs gene-disease associations for Orphanet-classified rare disorders, and DisGeNET, which documents gene-drug interaction records) through a reproducible three-stage gene symbol harmonization pipeline comprising HGNC identifier standardization, MyGene.info API mapping, and RapidFuzz fuzzy string matching. The resulting heterogeneous tripartite knowledge network encompasses 15,454 nodes and 35,131 edges, covering 2,249 clinically distinct rare diseases. A Graph Attention Network (GAT) is trained on this network using node-type classification as a pretext task, enabling the model to learn biologically informed node representations encoding the structural co-association of disease, gene, and drug entities. These representations are used to rank drug candidates for a query rare disorder via cosine similarity in the embedding space. The framework is explicitly positioned as a decision support tool for prioritizing existing gene-bridged drug-disorder connections rather than predicting novel associations. Held-out evaluation across 200 disorders demonstrates Hits@10 = 0.400 compared to a random baseline under 0.001, a greater than 400-fold improvement. The rank-4 retrieval of NITISINONE, the approved standard of care for hereditary tyrosinaemia type 1, for a query disorder sharing the HPD tyrosine catabolism pathway, without pathway annotations provided to the model, supports the biological plausibility of the learned embeddings. The complete pipeline is reproducible and is deployed as an interactive decision support interface on HuggingFace Spaces.

## 1 Introduction

Rare diseases affect over 300 million individuals worldwide yet more than 90% lack approved treatments (Global Genes 2023). Conventional drug discovery is economically prohibitive for small patient populations, making computational approaches to candidate identification an important research direction (Oprea et al. 2018; Pushpakom et al. 2019). Biomedical knowledge graphs, encoding genes, diseases, and drugs as nodes with documented associations as edges, provide a natural framework for this task.

The gene entity occupies a strategic position in this network: it serves as a molecular bridge between disease etiology and pharmacological action, enabling inference of candidate disease-drug connections through shared gene associations. Orphadata (Orphanet 2023) catalogs gene-isease associations for Orphanet-classified rare disorders; DisGeNET (Pinero et al. 2020) documents gene-drug interactions at scale. Used together through a gene bridge, these databases can link rare disorders to potential drug candidates. The practical barrier is that the two databases use incompatible gene naming conventions, preventing direct integration without explicit harmonization.

This paper presents a reproducible three-stage harmonization pipeline that resolves this barrier, yielding a heterogeneous tripartite knowledge network covering 2,249 rare diseases across 15,454 nodes and 35,131 edges. A Graph Attention Network (Velickovic et al. 2018) trained on this network using node-type classification as a pretext task (Hu et al. 2020) produces embeddings used for cosine similarity-based drug candidate ranking. The framework is designed as a decision support tool, ranking existing gene-bridged connections, not predicting novel ones, and is deployed as an accessible public interface.

Portions of the text in this manuscript were drafted with AI writing assistance (Anthropic Claude). All experimental design, data analysis, and scientific conclusions are the author’s own.

## 2 Related work

Sadeghi et al. (2022) developed a HinSAGE-based heterogeneous GNN for multi-label drug repurposing without rare-disease focus or attention mechanisms. He et al. (2024) proposed DRTerHGAT, a ternary heterogeneous GAT for Alzheimer’s drug repurposing, domain-limited and without cross-database harmonization. Huang et al. (2023) presented TXGNN, applying geometric deep learning across 17,080 diseases, but without addressing the Orphanet-DisGeNET harmonization challenge or providing a clinical deployment interface. The present work is distinguished by its harmonization pipeline as a primary contribution, its focus on the full Orphanet rare disease catalog, and its honest framing of ranking vs. novel prediction.

## 3 Materials and methods

### 3.1 Data sources

Orphadata XML parsing extracted 8,164 disorder-gene pairs spanning 3,961 rare disorders and 4,460 gene symbols. DisGeNET CSV provided 81,314 usable gene-drug records after dropping records with null gene or drug name fields. The framework pipeline is illustrated in Figure 1.

**Figure 1.**
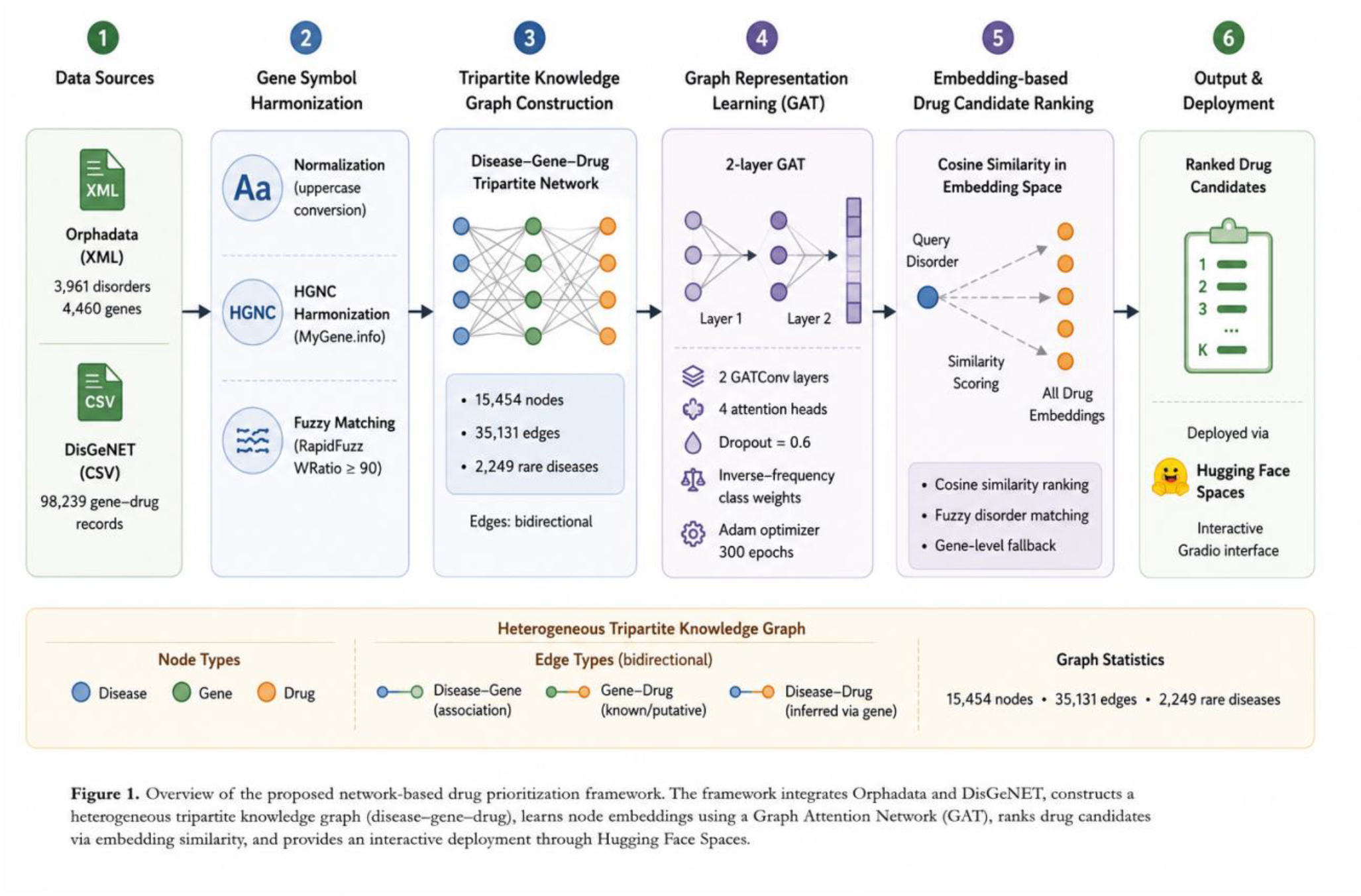
Overview of the proposed network-based drug candidate prioritization framework. The pipeline integrates Orphadata and DisGeNET through three-stage gene symbol harmonization, constructs a heterogeneous tripartite knowledge network, trains a Graph Attention Network as a pretext task, ranks drug candidates via embedding cosine similarity, and deploys through an interactive HuggingFace Spaces interface.

### 3.2 Gene symbol harmonization pipeline

Three sequential stages resolved nomenclature incompatibilities between the two databases. Stage 1 : Normalization: All gene symbols were uppercased and whitespace-stripped.

Stage 2 : HGNC mapping: Gene symbols were mapped to HGNC identifiers via the MyGene.info API (Xin et al. 2016), with EntrezGene IDs as fallback.

Stage 3 : Fuzzy matching: Residual mismatches were resolved using RapidFuzz WRatio matching at threshold 90 (Bachmann 2021).

Datasets were merged by inner join on standardized gene symbol, yielding 131,564 records before edge deduplication. The interaction_type field is populated for 28.1% of merged records, reflecting its availability only for DGIdb-sourced DisGeNET entries; this is reported transparently.

### 3.3 Heterogeneous tripartite knowledge network

A heterogeneous tripartite graph G = (V, E) was constructed with disorder, gene, and drug node types; bidirectional disorder-gene and gene-drug edges; and one-hot identity node features. Table 1 gives network statistics.

**Table 1.** Knowledge network statistics.

| Entity type | Count | Source database |
| --- | --- | --- |
| Disorder nodes | 2,249 | Orphadata (post-harmonization) |
| Gene nodes | 1,751 | Orphadata ∩ DisGeNET |
| Drug nodes | 11,454 | DisGeNET (post-harmonization) |
| Total nodes | 15,454 | - |
| Disorder-gene edges | 3,798 | Orphadata |
| Gene-drug edges | 31,333 | DisGeNET |
| Total edges (bidirectional) | 70,262 | - |

### 3.4 GAT architecture and training

A two-layer GAT (Velickovic et al. 2018) was implemented in PyTorch Geometric. Layer 1: GATConv to hidden dimension 8 with 4 attention heads (effective dimension 32), ELU activation, dropout p = 0.6. Layer 2: GATConv with 1 head, 3-dimensional logits, dropout. Adam optimizer (lr = 0.005, weight decay = 5e-4), 300 epochs over a stratified 70/15/15 node split.

The node distribution is structurally imbalanced: 74.1% drug, 14.6% disorder, 11.3% gene. Inverse-frequency class weights (w_disorder = 1.209, w_gene = 1.553, w_drug = 0.237) were applied to the cross-entropy loss. Node-type classification is the pretext task (Hu et al. 2020); the objective is biologically informed embeddings used for downstream drug candidate ranking.

### 3.5 Drug candidate ranking and deployment interface

After training, first-layer GAT embeddings are extracted. A query disorder is matched to the closest network node via RapidFuzz fuzzy matching, and cosine similarity between the disorder embedding and all 11,454 drug embeddings produces the ranked candidate list. All ranked drugs are connected to the query disorder through documented gene bridges. The system prioritizes existing connections, not novel predictions.

For disorders absent from the network, optional HGNC gene identifier input enables genelevel retrieval via averaged gene node embeddings. A Gradio interface is permanently deployed on HuggingFace Spaces (Ramani 2026a), accessible without specialist computational expertise. The complete implementation is available as a public Colab notebook (Ramani 2026b).

## 4 Results

### 4.1 Node-type classification

Table 2 reports per-class results on the held-out test set (n = 2,319 nodes).

**Table 2.** Per-class node classification results (n = 2,319)

| Node class | Precision | Recall | F1-score | Support |
| --- | --- | --- | --- | --- |
| Disorder | 0.332 | 0.580 | 0.422 | 338 |
| Gene | 0.663 | 0.721 | 0.691 | 262 |
| Drug | 0.920 | 0.773 | 0.840 | 1,719 |
| Macro average | 0.639 | 0.691 | 0.651 | 2,319 |
| Weighted average | 0.806 | 0.739 | 0.763 | 2,319 |

Macro F1 = 0.651 vs. unweighted majority-class baseline ≈ 0.28, a delta of 0.371 attributable to inverse-frequency weighting. Drug nodes achieve F1 = 0.840; gene nodes F1 = 0.691 despite minority class status; disorder nodes F1 = 0.422 (recall = 0.580), reflecting boundary ambiguity from shared neighbourhood structure with gene nodes. Figure 2 shows training dynamics.

**Figure 2.**
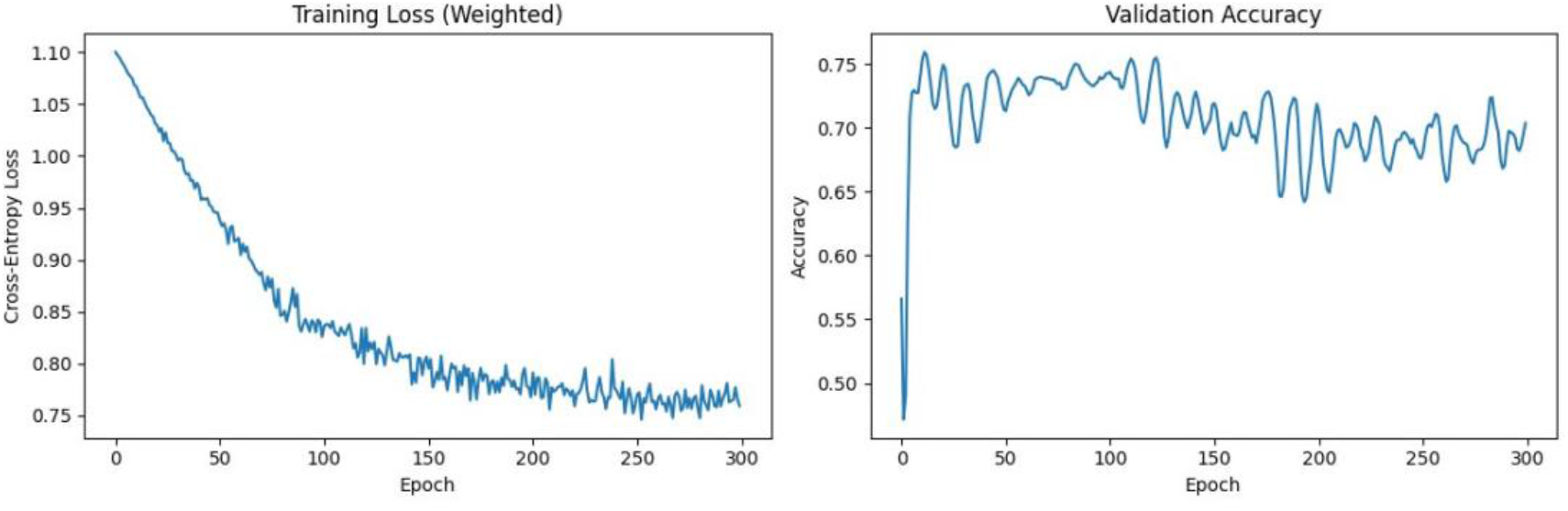
Training dynamics over 300 epochs. Left: weighted cross-entropy loss decreasing from 1.10 to 0.76. Right: validation accuracy stabilizing in the 0.65-0.75 range. Oscillation is attributable to full-batch dropout on the imbalanced node set.

### 4.2 Evaluation stability

Table 3 reports metrics across five stratified test partitions using the fixed trained model.

**Table 3.** Evaluation stability across five stratified test partitions.

| Partition | Accuracy | Macro F1 | ROC-AUC |
| --- | --- | --- | --- |
| 1 | 0.794 | 0.723 | 0.927 |
| 2 | 0.793 | 0.731 | 0.928 |
| 3 | 0.785 | 0.727 | 0.929 |
| 4 | 0.798 | 0.737 | 0.932 |
| 5 | 0.805 | 0.743 | 0.933 |
| Mean ± SD | 0.795 ± 0.007 | 0.732 ± 0.007 | 0.930 ± 0.002 |

Standard deviation does not exceed 0.007 across any metric, confirming performance stability.

### 4.3 Drug candidate ranking evaluation

Table 4 reports held-out ranking results across 200 disorders (20% drug mask per disorder).

**Table 4.** Drug candidate ranking evaluation across 200 disorders.

| Metric | K = 10 | K = 50 | Random baseline (K = 10) |
| --- | --- | --- | --- |
| Hits@K | 0.400 | 0.510 | < 0.001 |
| Recall@K | 0.153 | 0.231 | — |
| Mean reciprocal rank | 0.029 | — | — |

Hits@10 = 0.400 represents a greater than 400-fold improvement over the random baseline across 11,454 drug candidates. MRR = 0.029 reflects the challenge of top-1 recovery for pretext-task embeddings; retrieval-specific training is identified as a primary direction for future work.

### 4.4 Biological plausibility: Hawkinsinuria case study

Table 5 shows top-10 candidates for Hawkinsinuria (HPD mutations; tyrosine catabolism). The original query ‘Aspartylglucosaminuria’ was absent from the network (AGA gene lacks DisGeNET coverage); fuzzy matching identified Hawkinsinuria as the closest structural neighbour, demonstrating robustness to coverage gaps.

**Table 5.** Top-10 drug candidates for Hawkinsinuria (all connected via HPD gene bridge)

| Rank | Drug candidate | Cosine similarity | HPD gene bridge? |
| --- | --- | --- | --- |
| 1 | GNE-7883 | 0.9000 | Yes |
| 2 | RANOLAZINE | 0.8734 | Yes |
| 3 | DSP-2230 | 0.8686 | Yes |
| 4 | NITISINONE | 0.8558 | Yes (approved HT1 treatment) |
| 5 | AFACIFENACIN | 0.8412 | Yes |
| 6 | ADENOPHOSTIN A | 0.8404 | Yes |
| 7 | ZD-6126 | 0.8385 | Yes |
| 8 | BENOXINATE | 0.8378 | Yes |
| 9 | CX-5279 | 0.8336 | Yes |
| 10 | SERALUTINIB | 0.8319 | Yes |

NITISINONE ranks 4th. It is an FDA-approved HPPD inhibitor and standard of care for hereditary tyrosinaemia type 1, which shares the HPD tyrosine catabolism pathway with Hawkinsinuria. The model had no pathway annotations or approval labels; its rank-4 position among 11,454 HPD-connected candidates provides biological plausibility evidence for the learned embeddings.

## 5 Discussion

The central contribution is a reproducible harmonization pipeline making combined use of Orphadata and DisGeNET practically feasible for rare disease research. The resulting knowledge network is the largest heterogeneous integration of these two databases reported to our knowledge. From a health informatics perspective, the framework provides a biologically informed shortlist from what would otherwise be an unordered list of hundreds of gene-bridged candidates per disorder . Hits@10 = 0.400 means that for 40% of queried disorders, a known associated drug appears in the top 10 from a pool of 11,454 candidates.

The gap between Hits@10 = 0.400 and MRR = 0.029 is the main open problem. The embeddings are good enough for top-10 recovery in many cases but not for consistent top-1 placement. This is expected for pretext-task embeddings not specifically optimized for retrieval. A retrieval-specific training objective (contrastive learning over known disorder-drug pairs or ranking losses over held-out associations) is the most direct path to improvement.

43.2% of Orphadata disorders are absent from the network due to missing DisGeNET gene-drug coverage. This is a data gap, not a modeling gap, and reflects the current state of the rare disease pharmacological literature. One-hot node features carry no biological content; protein sequence embeddings or molecular fingerprints would be expected to improve both classification and ranking substantially. Full k-fold retraining was not feasible at this graph scale; mini-batch training strategies would enable this in future work.

The intended users are biomedical researchers and pharmaceutical scientists investigating rare diseases with limited existing drug options. The framework is appropriate as a first-pass computational screen to generate hypotheses for further experimental investigation, not as a clinical decision-making tool. Downstream experimental validation remains a necessary step before any clinical application.

## 6 Conclusion

This study presents a network-based framework for rare disease drug candidate prioritization, integrating Orphadata and DisGeNET through a reproducible three-stage harmonization pipeline into a heterogeneous tripartite knowledge network of 15,454 nodes and 35,131 edges covering 2,249 rare diseases. A GAT trained on node-type classification as a pretext task produces embeddings used for cosine similarity-based drug ranking. Held-out evaluation demonstrates Hits@10 = 0.400 across 200 disorders, more than 400 times better than random. NITISINONE’s rank-4 recovery for a tyrosine pathway disorder without pathway annotations provides biological plausibility evidence. The framework is reproducible, permanently deployed, and designed as an accessible decision support tool for rare disease drug candidate investigation.

## Data and code availability

Orphadata is publicly available at https://www.orphadata.com/. DisGeNET is publicly available at https://disgenet.com/. The complete implementation is available at https://colab.research.google.com/drive/19lbXIggc4ECxzA8bgeSuKeR6gBJxABzV?usp=sharing. A live deployment is available at https://huggingface.co/spaces/DiyaR2002/drug-repurposing.

## Acknowledgments

The author acknowledges the Orphanet consortium and the DisGeNET team for providing publicly accessible biomedical databases, and the open-source communities behind PyTorch Geometric, RapidFuzz, MyGene.info, and Gradio. The author used Claude (Anthropic) for manuscript drafting and code debugging. All experimental design, data analysis, and scientific conclusions are the author’s own.

## References

Bachmann M (2021) RapidFuzz: a fast string matching library for Python. J Open Source Softw 6(64):3030

Global Genes (2023) RARE facts. https://globalgenes.org/rare-facts/. Accessed 1 June 2025

He H, Zhu X, Chen M et al (2024) DRTERHGAT: a drug repurposing method based on the ternary heterogeneous graph attention network. J Mol Graph Model 130:108783

Hu W, Liu B, Gomes J et al (2020) Strategies for pre-training graph neural networks. In: Proc ICLR. arXiv:1905.12265

Huang K, Chandak P, Wang Q et al (2023) A foundation model for clinician-centered drug repurposing. medRxiv. 10.1101/2023.03.19.23287458

Oprea TI, Bologa CG, Brunak S et al (2018) Unexplored therapeutic opportunities in the human genome. Nat Rev Drug Discov 17(5):317–332

Orphanet (2023) Orphadata: free access data from Orphanet. https://www.orphadata.com/. Accessed 1 June 2025

Pinero J, Ramirez-Anguita JM, Sauch-Pitarch J et al (2020) The DisGeNET knowledge platform for disease genomics: 2019 update. Nucleic Acids Res 48(D1):D845–D855

Pushpakom S, Iorio F, Eyers PA et al (2019) Drug repurposing: progress, challenges and recommendations. Nat Rev Drug Discov 18(1):41–58

Ramani D (2026a) Rare disease drug candidate prioritization — live demo. Hugging Face Spaces. https://huggingface.co/spaces/DiyaR2002/drug-repurposing

Ramani D (2026b) Drug repurposing rare disease GAT pipeline. Google Colab. https://colab.research.google.com/drive/19lbXIggc4ECxzA8bgeSuKeR6gBJxABzV?usp=sharing

Sadeghi S, Lu J, Ngom A (2022) An integrative heterogeneous graph neural network-based method for multi-labeled drug repurposing. Front Pharmacol 13:908549

Velickovic P, Cucurull G, Casanova A et al (2018) Graph attention networks. In: Proc ICLR. arXiv:1710.10903

Xin J, Mark A, Afrasiabi C et al (2016) High-performance web services for querying gene and variant annotation. Genome Biol 17(1):91

